# Neural indicators of human gut feelings

**DOI:** 10.1101/2021.02.11.430867

**Authors:** Ahmad Mayeli, Obada Al Zoubi, Evan J. White, Sheridan Chappelle, Rayus Kuplicki, Ryan Smith, Justin S. Feinstein, Jerzy Bodurka, Martin P. Paulus, Sahib S. Khalsa

## Abstract

Understanding the neural processes governing the human gut-brain connection has been challenging due to the inaccessibility of the body’s interior. Here, we investigated neural responses to gastrointestinal sensation using a minimally invasive mechanosensory probe by quantifying brain, stomach, and perceptual responses following the ingestion of a vibrating capsule. Participants successfully perceived capsule stimulation under two vibration conditions (normal and enhanced), as evidenced by above chance accuracy scores. Perceptual accuracy improved significantly during the enhanced relative to normal stimulation, which was associated with a more rapid detection of the stimulation and reduced reaction time variability. Stomach stimulation induced early and late neural responses in parieto-occipital leads near the midline. Moreover, these ‘gastric evoked potentials’ showed intensity-dependent increases in amplitude and were significantly correlated with perceptual accuracy. These findings highlight a unique form of enterically-focused sensory monitoring within the human brain, with implications for understanding gut-brain interactions in healthy and clinical populations.

## Introduction

The human brain must decipher a multitude of complex signals originating from within the body in order to maintain homeostasis and optimally sustain life. Interoception, the process by which the nervous system senses, interprets, and integrates signals originating from within the body across conscious and nonconscious levels (1), is a vital component of this homeostatic machinery maintaining internal stability in the face of changing environments. However, interoception remains poorly understood despite both growing scientific interest (2–5) and the recognition that certain psychiatric (6) and neurological (7, 8) disorders may manifest through abnormal neural processing of interoceptive signals.

Most research on the gut-brain connection has focused on the cellular and molecular mechanisms of afferent interoceptive signal transmission in nonhuman animals (9). For example, in the alimentary tract, rapid cell-specific peripheral sensors of osmotic balance (10), glucose (11, 12), and mechanical stretch (13, 14) have been identified that enable organisms to quickly regulate feeding/drinking behaviors before the onset of relevant blood-level changes. Neurons in the insular cortex have been identified to play a key role in the integration of these viscerosensory signals and the adaptive estimation of upcoming needs (15, 16) – providing a focal point within the central nervous system for understanding interoceptive predictive processing. However, the brain also exerts powerful influences on the gut and multiple descending pathways linking the stomach and brain have been identified. For example, the insular and medial prefrontal cortices send parasympathetic projections to the stomach through the vagus nerve, whereas the primary motor and somatosensory cortices send sympathetic projections to the stomach through spinal efferents (17).

In humans, the lack of appropriate techniques for easily accessing the body’s interior has hindered the study of interoceptive processing. Some of these limitations have begun to be addressed through the development of pharmacological or mechanosensory perturbations of cardiac and respiratory sensation (18, 19), yet there remains a dearth of minimally invasive probes for assessing other organ systems such as the gut. Most prior studies evaluating the conscious perception of gastrointestinal sensations have used invasive approaches involving the insertion of inflatable balloons or electrical probes into the esophagus (20, 21), stomach (22, 23), colon (24), or rectum (25). While such approaches have shown the ability to engage putative interoceptive cortical circuitry (i.e., insular and somatosensory cortices)(26) and have revealed insights into the gut-brain axis in gastrointestinal disorders (27, 28), the invasiveness of these approaches has made it challenging to conduct such research in human participants.

In the current study, we developed a minimally invasive probe targeting perceptions of the gastrointestinal system via ingestion of a vibrating capsule. Using a design inspired by signal detection theory, we combined the mechanosensory stimulation of stomach signals with perceptual measurement of stomach sensations and continuous recording of electroencephalogram (EEG), electrogastrogram (EGG), and other peripheral physiological signals. We identify (1) reliable signatures of gastrointestinal perception at the individual subject level and (2) differential effects in the brain based on the degree of stimulation, as measured by evoked response potentials (ERP) using EEG. We suggest that this minimally invasive approach could serve as a useful method for understanding gut-brain interactions across a variety of human health conditions.

## Results

40 healthy human participants (19 female, average age = 22.90 ± 4.56 years, range between 19 and 39 years, average BMI = 24.18 ± 3.03, range between 18.24 and 32.28) completed the study and met quality assurance criteria for inclusion in the analysis.

### Vibratory Stomach Stimulation Modulates Gut Sensation

We examined whether the noninvasive delivery of vibratory stimulation to the stomach would yield reliable signatures of gastrointestinal perception at the individual subject level. Following ingestion of the capsule, participants were instructed to press a button anytime they consciously perceived a feeling in their gut (Figure 1).

**Figure 1:**
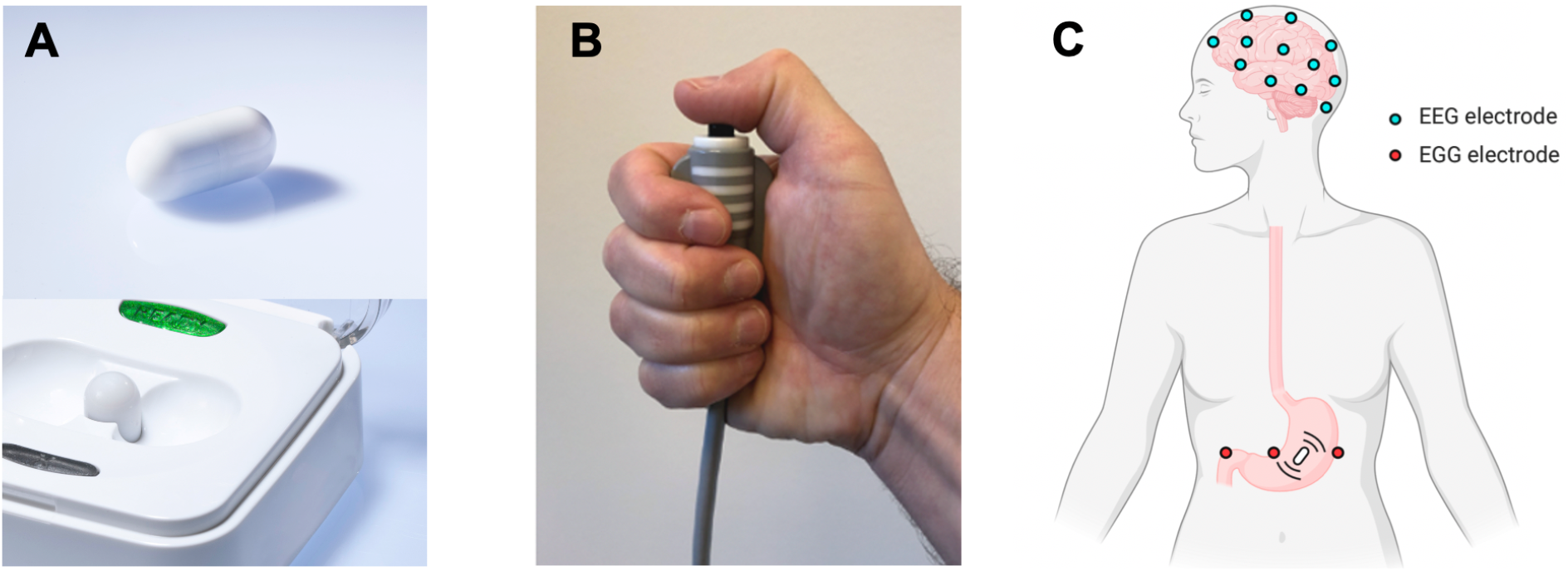
**A)** Vibrating capsule and activation base. **B)** The push button which participants were asked to press each time they detected a capsule-induced stomach sensation. **C)** Scalp electroencephalogram (EEG) and stomach electrogastrogram (EGG) lead placement.

The analysis of button-press responses showed that participants could successfully detect vibration stimuli under both normal and enhanced stimulation conditions, Normalized A prime (mean ± standard deviation [STD]) = 2.49 ± 0.40 and 2.84 ± 0.25, respectively. Perceptual accuracy increased significantly under enhanced versus normal stimulation (paired t-test: *p* = 4.11e-08, *t(39)* = 6.79, Cohen’s *d* = 1.1) (Figure 2A). Response latencies (time from capsule stimulation to button press) during normal and enhanced blocks were as follows: average response latency (mean ± STD) = 1.06 ± 0.33 seconds (normal stimulation) and 0.74 ± 0.23 seconds (enhanced stimulation); STD of response latency (mean ± STD) = 0.46 ± 0.14 (normal stimulation) and 0.28 ± 0.13 seconds (enhanced stimulation). Response latencies decreased significantly under enhanced versus normal stimulation for both the average response latency (paired t-test: *p* = 7.61e-07, and *t(38)* = 5.91, Cohen’s *d* = 1.0) and for the STD of response latency (paired t-test: *p* = 3.20e-08, and *t(38)* = 6.92, Cohen’s *d* = 1.1) (Figure 2B and C). One participant did not respond to any of the normal vibration stimuli; therefore, for comparisons of response latency/STD between normal and enhanced conditions, 39 participants were included. The binomial test showed that all 40 participants performed significantly above chance (*p* < 0.05) during enhanced stimulation, whereas 5 participants performed below chance (*p* ≥ 0.05) during normal stimulation.

**Figure 2:**
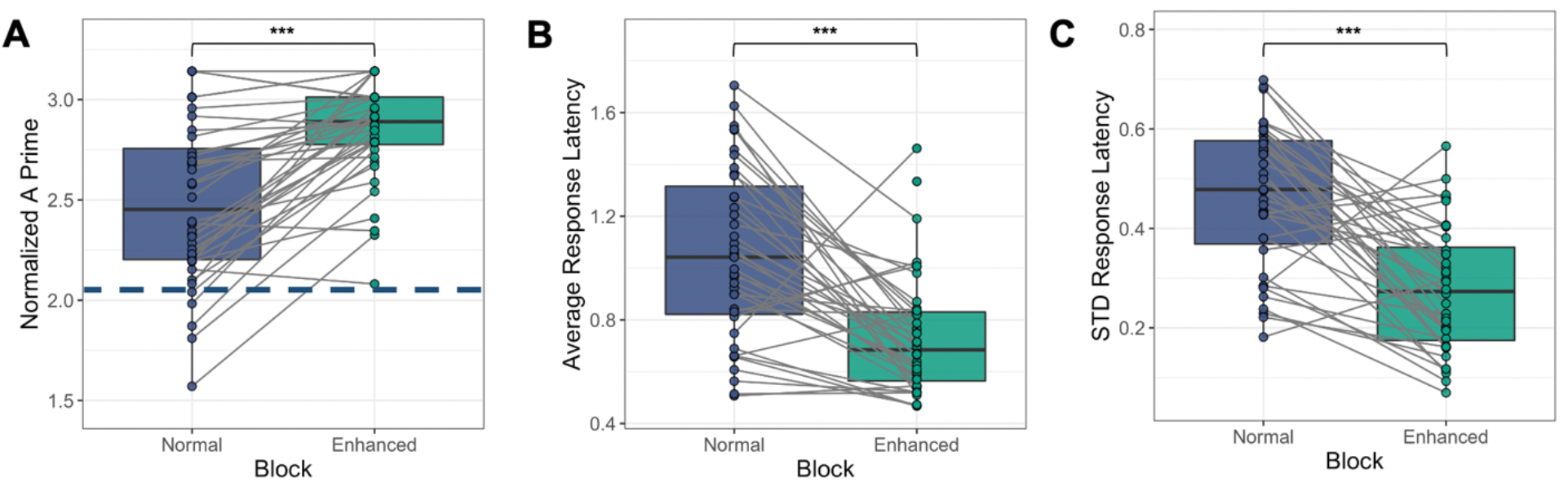
Perceptual accuracy measures during the normal and enhanced stimulation blocks based on button-presses signifying gut feelings during vibratory stimulation of the stomach. Gray lines present changes in individual performance from the normal to enhanced block. **A)** Normalized A prime (dashed line shows chance performance based on binomial expansion); **B)** Average response latency (in seconds); and **C)** Standard deviation (STD) of the response latency (in seconds). The participants’ performance increased significantly with enhanced stimulation across all three measures, indicating the paradigm effectively induced changes in gut sensation. \*\*\**P* < 0.001.

### Sex Effects Do Not Account for Observed Changes in Gut Sensation

Repeated measures analysis of variances (ANOVAs) of the normalized A prime, average response latency, and the STD of response latency as dependent variables (applied separately for each variable), with sex and block as fixed factors showed no significant differences between the males and females for any of these dependent factors, despite significant differences between normal and enhanced stimulation for all dependent variables (Table S1).

### Parieto-occipital ERP Indicators of Gut Sensation

Early negative and late positive deflections in the ERP signal emerged around 100 milliseconds and 400 milliseconds (ms) after stimulation onset, respectively. The early negative deflections peaked around 150 ms whereas the late positive deflections peaked around 600 ms and lasted up to 3000 ms (i.e., the duration of the vibration stimulus). These changes were maximally located over the midline centro-parietal electrode sites for both conditions. Additionally, inspection of the signal showed intensity-dependent differences in early negative and late positive deflections.

For the late response period a significant main effect of condition, *F(1)* = 19.48, *p* < 0.001, *η2* = 0.34, revealed that the enhanced stimulation condition (Mean (*M)* = 3.93 *microvolts (μV)*) resulted in a larger ERP amplitude than the normal stimulation condition (*M* = 2.48 *μV*) for the nine channels showing the largest responses (CP1, CP2, P3, Pz, P4, POz, O1, Oz, and O2). There also was a main effect of Channel, *F(1.85)* = 10.13, *p* < 0.001, *η2* = 0.21; however, the two-way interaction between Channel and Condition, *F(2.51)* = 2.84, *p* = 0.074, was non-significant. Figure 3A displays the averaged ERP waveform for the nine channels showing the largest responses, and Figures 3B and 3C show the scalp topography maps for early and late deflections in each condition. Thus, the enhanced condition resulted in a larger late ERP amplitude than the normal condition, and this enhancement was not substantially modulated by the measurement site (Table S2, S3 and Figure S1). The repeated measures ANOVA applied to the averaged ERPs among midline centro-parietal electrodes between 400 and 700 ms showed no significant sex differences between males and females, *F(1)* = 3.405, *p* = 0.069 and no significant interaction between Sex and Condition, *F(1)* = 0.063, *p* = 0.803, but a significant difference by Condition, *F(1)* = 9.044, *p* = 0.004.

**Figure 3:**
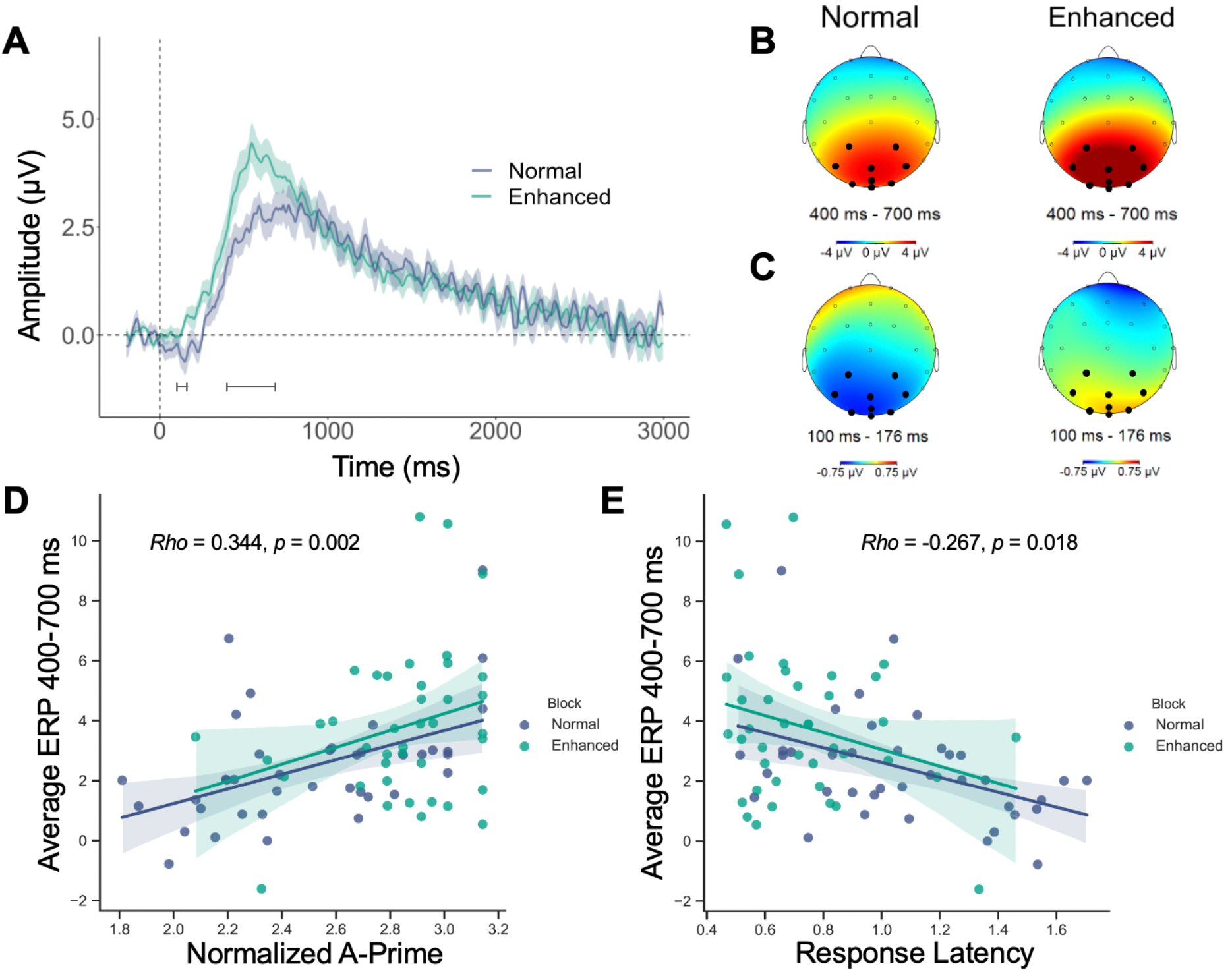
Parieto-occipital event related potential (ERP) indicators of gut sensation and their association with perceptual accuracy measures during normal and enhanced stimulation. **A)** The average ERP waveforms during the normal (blue) and enhanced (green) blocks for channels CP1, CP2, P3, Pz, P4, POz, O1, Oz, and O2 (indicated as black dots in the B) and C) scalp maps). Intensity-dependent differences were examined during early (e.g. 100 to 176 ms) and late (e.g. 400 to 700 ms) windows (marked with horizontal brackets). Shaded areas represent the standard error of the mean for the ERP signal at each time point. Time-zero represents the earliest onset of vibratory stimulation. The presented waveforms were calculated from the average mastoid-referenced EEG. **B)** Scalp topography during the late window after the vibration onset relative to the pre-stimulus baseline for the normal and enhanced conditions. **C)** Scalp topography during the early window after the vibration onset relative to the pre-stimulus baseline for the normal and enhanced conditions. **D)** The positive association between the late ERP signal strength (averaged signal among CP1, CP2, P3, Pz, P4, POz, O1, Oz, and O2 channels) and perceptual accuracy (normalized A prime) was significant across both conditions. **E)** The negative association between late ERP signal strength (averaged signal among CP1, CP2, P3, Pz, P4, POz, O1, Oz, and O2 channels) and response latency was significant across both conditions. For plots D and E, the Rho and p values represent the Spearman partial correlation results controlling for condition. Shaded areas correspond to the 95% confidence interval for the regressions.

An early negative waveform peaking around 150 ms was also observed across the averaged electrodes during the normal but not enhanced condition. The repeated measures ANOVA applied to the averaged waveform among midline centro-parietal electrodes between 100 and 176 ms (inclusive of the 150 ms peak) showed no significant sex differences between males and females, *F(1)* = 0.135, *p* = 0.714; no significant interaction between Sex and Condition, *F(1)* = 0.171, *p* = 0.680; but a significant difference by Condition, *F(1)* = 9.111, *p* = 0.003.

To evaluate the potential role of motor activity (i.e., button presses) in the observed ERP signals, in a separate analysis we examined early and late ERP responses to “false positive” button presses. There were no clear ERP components observed when participants responded to non-vibration stimuli (false positive button presses) or when they missed responding to the vibration (false negative) (Figures S2 and S3 in supplement).

### Neural Gut Sensation Indicators Relate to Perceptual Accuracy

There was an association between the average LPP amplitude and perceptual accuracy (as measured by normalized A prime) after controlling for Condition (i.e., normal/enhanced), *Rho* = 0.344, *p* = 0.002 (Figure 3D). In comparison, there was a negative association between LPP and response latency, *Rho* = −0.267, *p* = 0.018 (Figure 3E) but no association between LPP and the average STD of response latency, *Rho* = −0.196, *p* = 0.086.

### Vibratory Stomach Stimulation Does Not Modulate EGG Signals

The repeated measures ANOVA examining the EGG total power showed no significant differences between the baseline, normal, and enhanced blocks, *F(2, 78)* = 2.50, *p* = 0.089, *η2* = 0.062 (Figure 4). To evaluate potential frequency-specific changes in the EGG signal, we conducted a separate repeated measures ANOVA for the three main frequency ranges (bradygastria [0.5 to 2.5 cycles per minute, cpm], normogastria [2.5 to 3.5 cpm], and tachygastria [3.75 to 9.75 cpm]) as a function of the three Conditions (baseline, normal, enhanced). There was a significant main effect of frequency band, *F(1.25)* = 5.75, *p* = 0.015, *η2* = 0.13; however, no significant main effect of Condition was observed *F(1.99)* = 3.0, *p* = 0.056, *η2* = 0.07), and the two-way interaction between frequency bands and Condition was non-significant *F(2.17)* = 1.49, *p* = 0.230, *η2* = 0.038 (Figure S4 in supplement).

**Figure 4:**
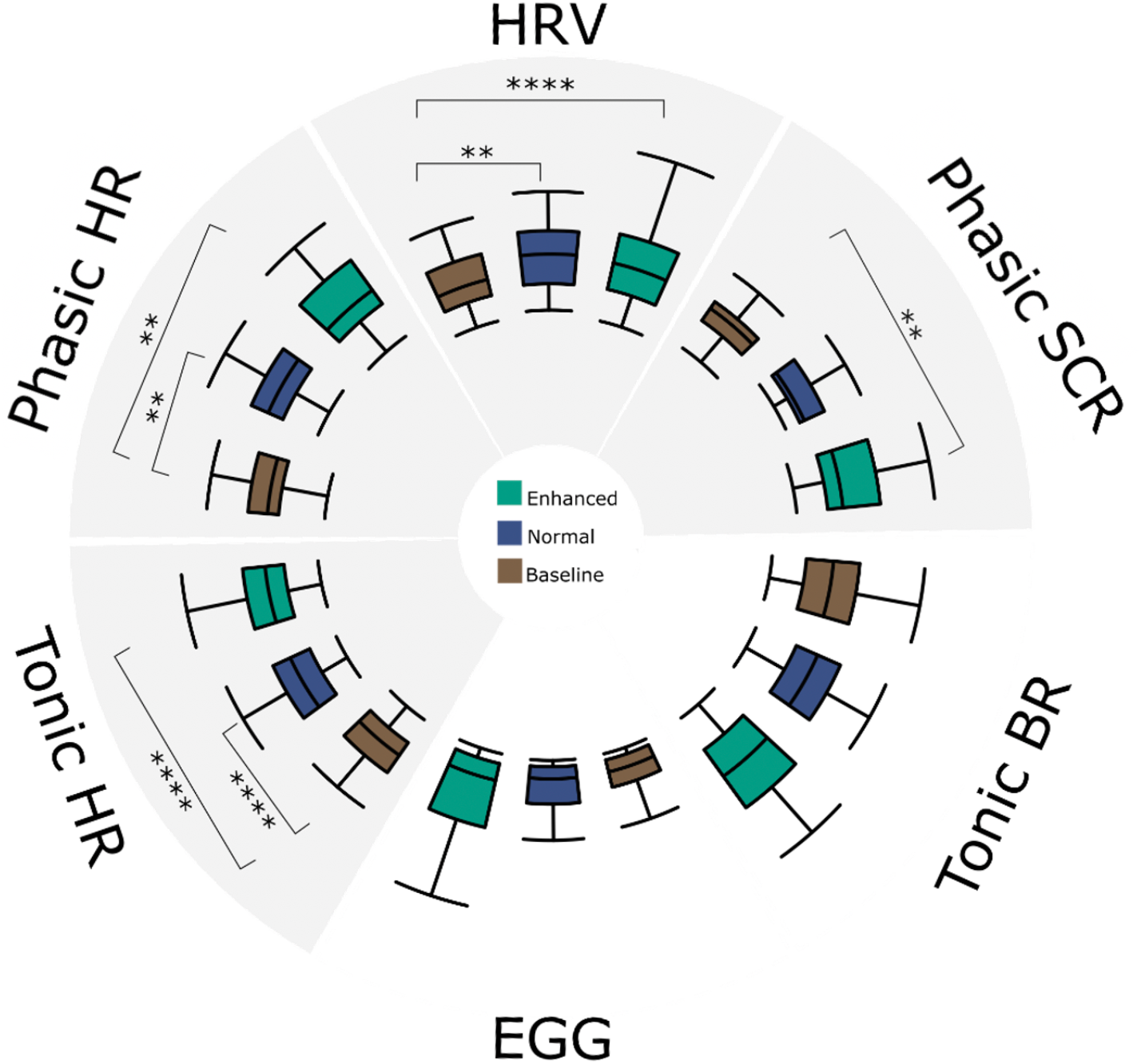
Influence of vibratory stomach stimulation on peripheral physiological measures during phasic (rapid – event related) and tonic (slow – entire block) periods. Shaded quadrants indicate signals with significant stimulation-associated changes from baseline, with brackets denoting post-hoc comparisons after Bonferroni-correction. **Top-right:** phasic skin conductance response (SCR) amplitudes differed from baseline during the enhanced stimulation condition only. **Top-left:** phasic heart rate (HR) responses differed from baseline during both the enhanced stimulation conditions. **Bottom-left:** tonic HR differed from baseline during both the normal and enhanced stimulation conditions. **Top:** tonic heart rate variability (HRV) differed from baseline during both the normal and enhanced stimulation conditions. **Bottom:** Total electrogastrogram (EGG) power across all physiologically relevant spectrums did not differ for each stimulation condition. **Bottom-right:** estimated breathing rate (BR) responses did not change across blocks. **P < 0.01, ****P < 0.0001.

### Vibratory Stomach Stimulation Modulates Peripheral Physiological Signals

#### Skin Conductance Responses (SCRs)

The maximum value of phasic activity differed significantly across Conditions, *F(2,78)* = 8.06, *p* < 0.001, *η2* = 0.12. Bonferroni adjusted post-hoc analyses revealed significant differences between the baseline and enhanced conditions (paired t-test: *t(39)* = −3.69, *p* < 0.01, Cohen’s *d* = −0.87); but no differences between the baseline and normal (paired t-test: *t(39)* = −2.00, p=0.16), nor between the normal and enhanced conditions (paired t-test: *t(39)*= −2.21, *p* = 0.33) (Figure 4).

#### Cardiac Responses

Tonic (i.e., basal) heart rate (HR) activity differed significantly across Conditions, *F(2,78)* = 46.5, *p* < 0.0001, η2 = 0.07. Bonferroni adjusted post-hoc analyses revealed significant differences between baseline and both normal (paired t-test: *t(39)* = −7.86, *p* < 0.00001, Cohen’s *d* = −0.60), and enhanced conditions (paired t-test: *t(39)* = −10.86, p < 0.00001, Cohen’s *d* = −0.56), but not between the normal and enhanced conditions (paired t-test: *t(39)* = −1.31, *p* = 0.59) (Figure 4). Phasic HR activity (i.e., fast responses specific to the 3-second vibration period) differed significantly across Conditions, *F(2,72)* = 8.719, *p* < 0.001, *η2* = 0.12. Bonferroni adjusted post-hoc analyses revealed significant differences between baseline and normal (paired t-test: t(39) = −3.40, p < 0.01, Cohen’s *d* = −0.81) and enhanced conditions (paired t-test: *t(39)* = −3.72, *p* < 0.01, Cohen’s *d* = −0.80), while no differences between the normal and enhanced conditions (paired t-test: *t(39)* = −0.21, *p* = 1) (Figure 4). Heart rate variability (HRV), as measured through the SDNN metric, differed significantly across Conditions, *F(2,78)* = 14.23, *p* < 0.00001, *η2*= 0.043. Bonferroni adjusted post-hoc analyses revealed significant differences between baseline and the normal (paired t-test: *t(39*) = −3.88, *p* < 0.01, Cohen’s *d* = −0.49) and enhanced conditions (paired t-test: *t(39)* = −5.29, *p* < 0.00001, Cohen’s *d* = −0.40), while there were no differences between normal and enhanced (paired t-test: *t(39)* = −1.27, *p*=0.64) (Figure 4).

#### Respiratory Responses

Tonic breathing rate (BR) activity did not show any significant differences across blocks, *F(2,78)* = 1.09, *p* =0.34, *η2*= 0.005 (Figure 4). Descriptive statistics for all peripheral physiological signals are listed in Table S5.

### Vibratory Stomach Stimulation Induces Cross-Channel Respiratory Interference

The repeated measures analysis of interoceptive intensity ratings pre- and post- stimulation revealed significant differences between the rating types, *F(2.367)* = 45.214, *p* < 0.001, *η*2 = 0.537, the different time points, *F(1)* = 35.254, *p* < 0.001, *η*2 = *0.475*, and a significant interaction between time point and rating types, *F(2.51)* = 29.909, *p* < 0.001, *η2* = 0.434. Bonferroni-adjusted post-hoc comparison testing showed significant increases in both stomach/digestive and respiratory sensation intensity (paired t-test: *t(39)* = *9.861*, *p* < 0.001, Cohen’s *d* = 1.56 for stomach/digestive intensity, and *t(39)* = 2.782, *p* = 0.033, Cohen’s *d* = 0.44 for respiratory intensity). The estimated marginal means of pre- and post- stimulation for all four measures are presented in Table S6. The magnitude of the stomach sensation intensity rating changes was larger than that for respiratory sensation. There were no significant differences for heartbeat or muscle tension ratings (corrected *p-values* are 0.181 and 1, respectively). Figure 5 illustrates the sensation intensity ratings across the experiment. There were also significant correlations between changes in stomach/digestive, breath, and heartbeat intensity ratings (Post-Pre) as shown in supplementary table S7. However, there were no significant correlations between ratings of muscle intensity and the other organs. Finally, there were no significant differences between males and females observed for any of the interoceptive intensity ratings (Table S8).

**Figure 5:**
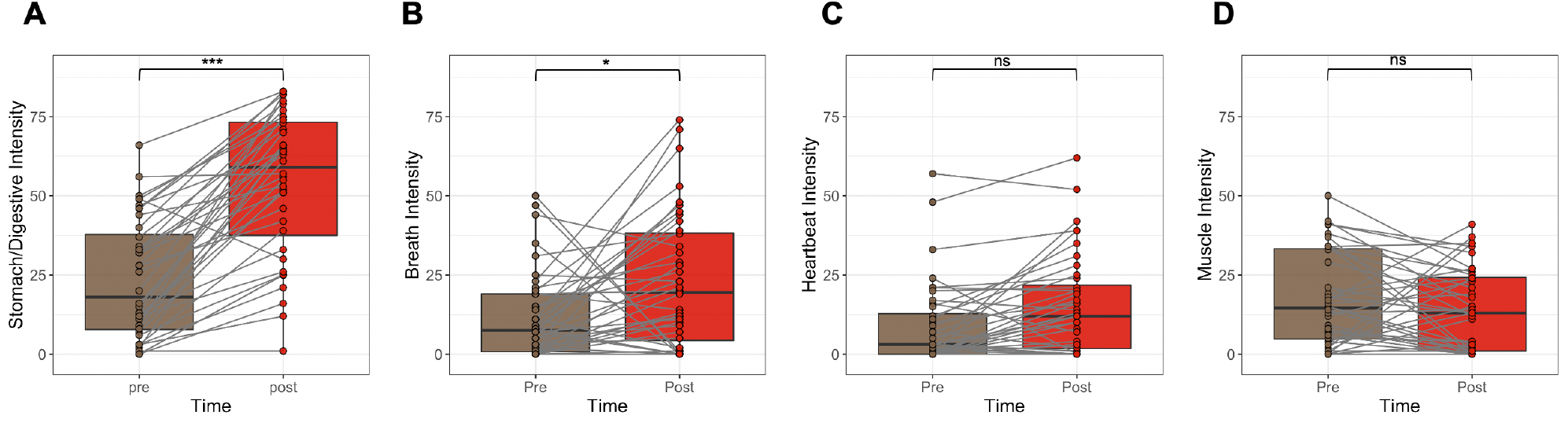
Self-reported intensity ratings of different interoceptive sensations experienced before (Pre) and during stimulation (Post; these ratings were provided retrospectively, and encompassed sensations experienced during both blocks). **A)** Stomach/Digestive, **B)** Breath, **C)** Heartbeat, and **D)** Muscle tension ratings. Stimulation-induced intensity ratings increased for both stomach and breath sensations. Gray lines show changes in ratings for each individual. \**P* < 0.05, \*\*\**P* < 0.001; ns, not significant.

## Discussion

In the current study we demonstrated that a minimally invasive form of mechanosensory gastrointestinal stimulation reliably changes the perception of gut feelings. Vibrating stimulation in the stomach induced evoked responses in the brain in midline parieto-occipital electrodes at both early and late periods following vibration of the capsule. These ‘gastric evoked potentials’ (GEPs) responded to stimulation and were significantly correlated with perceptual accuracy. Furthermore, our observations of intensity-dependent increases in both perceptual accuracy measures and GEP amplitudes between the normal and enhanced forms of stimulation indicate the reliability and effectiveness of this minimally invasive assay of gastrointestinal interoception. Mechanosensory stomach stimulation was not associated with the modulation of intrinsic gastric myoelectric rhythms, though it was accompanied by changes in peripheral (i.e., cardiac and electrodermal) indicators of arousal. Overall, these results highlight the presence of a distinct form of enterically-focused sensory monitoring within the human brain.

Prior functional neuroimaging investigations during mechanosensory balloon distension of the esophagus and colorectum showed activation in the primary/secondary somatosensory (SI/SII) and insular cortices (26–28). We found no ERP pattern consistent with activation in these areas, and instead observed midline parieto-occipital evoked potentials during mechanosensory stomach stimulation. However, these ERP findings concur with a recent study documenting the presence of a ‘gastric network’ in the brain (29). This gastric network included bilateral SI/SII nodes, but notably, a greater number of nodes were located in the posterior cingulate sulcus, superior parieto-occipital sulcus, dorsal precuneus, retrosplenial cortex, as well as the dorsal and ventral occipital cortex. In the study by Rebollo et al., it was predominantly the posterior midline brain regions that coactivated with the EGG signal under resting conditions, and different nodes within this network showed patterns of early and delayed changes in functional connectivity in relation to the EGG signal. Although 32-channel EEG measurements do not provide the spatial resolution necessary to optimally pinpoint the cerebral source of the observed parieto-occipital LPPs, the posterior midline brain regions observed by Rebollo et al. are located within a plausible set of brain regions for generating this result. Despite our focus on interoceptive awareness in the current study, the ERP results did not show patterns suggestive of an insular or somatosensory genesis. Instead, our findings highlight the possibility that posterior midline brain regions could play a role in interoceptive awareness for stomach sensations. This is consistent with another study based on magnetoencephalography showing an association between EGG signals and midline posterior parietal and occipital regions (30). Using a causal interaction analysis, Richter et al. found greater evidence for information transfer from the gut to the brain than the opposing direction (brain to gut). Taken together, our data support the hypothesis that gastric and intestinal interoception may be processed in posteromedial brain regions.

A recent landmark study found that deep posteromedial cortex stimulation was associated with an ‘out of body’ experience (i.e., dissociation) in an epileptic individual with intracranial recording electrodes implanted for seizure localization (31). In particular, the patient exhibited abnormal oscillatory activity in the deep posteromedial cortex prior to the onset of seizures associated with dissociation, and electrical stimulation of this region triggered the same oscillations and dissociation symptoms that mimicked the pre seizure aura (a similar pattern of findings was firmly established in the mouse using optogenetic stimulation of layer 5 retrosplenial neurons and induction of dissociation-like episodes using ketamine). The posterior cingulate cortex, which is adjacent to the retrosplenial cortex, is another region which has shown selective recruitment during stimulation of the gastric fundus to elicit fullness sensations as well as pain (32). This same region has been implicated in the hierarchical mapping of autonomic nervous system input via demonstrations of abnormal activity in this region in the setting of pure autonomic failure (33). Theoretical proposals have also pinpointed this region as a key substrate for the re-mapping of first-order body representations subserving emotion and conscious awareness in response to ongoing behavioral and environmental contexts (34). Collectively, these studies further emphasize the role of posterior midline structures in gastrointestinal interoception.

Neither group nor individual level analyses provided evidence that the vibrating capsule stimulation changed the frequency of stomach activity as indexed by the EGG signal. This result is important because it suggests that the afferent/ascending mechanosensory stimulation delivered by the capsule did not evoke a detectable visceromotor (regulatory) gut response, at least at the systems level of measurement employed here. Additional studies would be needed to establish the same finding in clinical populations, particularly for those in which gastric dysrhythmias are commonly encountered (35). However, our analysis of other physiological signals revealed evidence that capsule stimulation induced changes in peripheral indicators of arousal. Specifically, phasic vibration-associated changes in heart rate and electrodermal responses were superimposed upon a tonic pattern of heightened heart rates (and altered HRV) occurring throughout both capsule stimulation conditions. These physiological findings are consistent with a pattern of both acute and sustained arousal changes that were induced by capsule stimulation, with a maximial arousal shift occurring during the enhanced condition. Whether such arousal responses reflect sensory or regulatory responses to gut stimulation deserves further study, particularly given arguments that arousal events can bias the attentional prioritization of information in favor of top-down processing (36).

Minimally invasive measurement of gut sensation is an important advance given the difficulty of gaining access to the gastrointestinal system, and these finding open the possibility for future clinical disorders investigations. There are a number of poorly understood conditions in which abnormal gut sensation is a part of symptom-based care settings, including eating disorders (e.g., anorexia and bulimia nervosa)(37, 38), somatic symptom disorders (39, 40), functional neurological disorders (8, 41), and functional bowel disorders (e.g., irritable bowel syndrome, functional dyspepsia, or functional bloating (42, 43). Clinicians are often faced with a patient reporting prominent gut symptoms without a clear medical explanation, despite the understanding that these disorders involve abnormal nervous system representation of internal sensory information (6–8). The current approach enables examinations of the relationship between gastrointestinal symptoms, illness severity, GEPs, and accuracy for detecting stomach stimulation in these clinical populations. Based on the sensitivity of this measure, particularly the ability to derive robust perceptual estimates at the individual level, such studies could potentially contribute to a better accounting of the pathophysiological manifestation of symptoms (e.g., how do internally arousing stimuli contribute to the formation of the ‘symptom scaffold’, a process whereby bodily sensations are systematically interpreted as threatening? (41, 44)). Another promising future direction could be to combine this paradigm with more sophisticated computational modelling approaches, which is an area of interoception research that has been substantially limited by a lack of compatible tools (5, 6).

Studies using invasive approaches have previously identified an evoked potential during esophageal, duodenal, and anorectal stimulation (21, 45). This ‘visceral evoked potential’ was characterized by a triphasic response pattern (P1, N1, P2), usually resolving within 300 ms. It is important to note that we did observe a deflection in the normal condition consistent with the time course of N1 ellicited in other sensory pardigms (e.g., visual or auditory (46)); however the interpretation of this deflection is limited by overlap with residual alpha activity in the averaged waveforms and considering the negative shift begins near the onset of stimulation in some leads (Figure S1). Previous visceral evoked potential work also differs from the signal we observed, which consisted of a monophasic peak with a delayed onset starting around 200 ms after stimulation, peaking at 600 ms, and lasting up to 3000 ms centered over the Pz and POz electrode sites. There are several potential explanations for this discrepancy. First, the studies in question used discrete painful or aversive forms of stimulation (e.g., mechanical distension or electrical shock) to induce a sensory response, whereas capsule stimulation evoked protracted nonpainful and nonaversive sensations via vibration. Second, the aforementioned studies used a single recording channel via an electrode placed at the vertex (Cz), as opposed to the 32 channels recorded across the head in the current study. We are unable to speculate on whether these studies would have seen a stronger signal localized to more posterior parietal or occipital leads if they had included a larger array of recording electrodes. Third, the invasive studies involved stimulation under passive instruction conditions, whereas the current study paired a continuous top-down (i.e., goal-directed) focus of interoceptive attention with a sustained motor response (the button press and hold), a cognitive task which presumably recruits a broader set of cortical regions. This distinction is important when considering the neural processes potentially contributing to the different phases of observed neural responses. For example, motor reaction times to passive auditory, visual, and somatosensory stimulation are typically on the order of 120 to 150 ms in duration (47), whereas the average reaction times in the current study were about 1 second for the normal condition and 0.75 seconds for the enhanced condition (the vibration stimulus took about a quarter second to ramp up, Figure S5, which could have also impacted response times). While neural processing related to the motor response following perception of stomach stimulation likely contributed to some of the delays in the observed ERP signal, it is insufficient to completely explain it. Thus, the observed ERPs could potentially reflect a combination of sensory, cognitive, and motor activity, which might help to explain why the ERPs had extended response profiles. We use the term GEP for the observed signal in part for simplicity and to acknowledge the novelty of this deflection in response to gastrointestinal stimulation. The prolonged time course, morphology, and scalp distribution is similar to that of the LPP (48), which is generally thought to reflect reciprocal interactions between frontal and parieto-occipital regions, and which is sensitive to autonomic and self-reported indices of arousal (49) and emotion regulation strategies (48). The scalp distribution and onset of the GEP waveform could also be consistent with the P3b component, a marker of stimulus recognition (50), although the GEP observed herein demonstrates a more protracted positivity than typical P3b studies. In sum, vibratory gastrointestinal stimulation evoked GEPs characterized by a protracted positive deflection in the ERP waveform, and the amplitude of this deflection was modulated by stimulation intensity in a manner seemingly analogous to arousal modulation of visually evoked LPPs (49). The current findings are preliminary with respect to the functional interpretation, and empirical work is needed to more precisely delineate the perceptual and attentional functions that elicit and modulate its components.

This study has certain limitations. We cannot completely rule out the contribution of afferent respiratory stimulation to the observed posteromedial GEP signals given the increased respiratory sensation reports (for example, due to potential transmission of vibratory stimulation to the lungs through the respiratory diaphragm, which sits on top of the stomach). We would expect this degree of interference to be minimal, given the disproportionate magnitude of the effect on stomach sensation and lack of observed changes in respiratory rate. Another possibility is that attending to stomach sensations also promotes attention to respiration. One important limitation of scalp EEG recordings is that they cannot easily detect localized signals originating from deep within the cortex. Thus, we cannot conclusively exclude the possibility that the capsule stimulation also evoked changes in deep subcortical structures previously implicated in gut sensation such as the insular cortex, thalamus, and brainstem. Studies involving positron emission tomography (PET) scanning could overcome this limitation. Functional magnetic resonance imaging (fMRI) or magnetoencephalography (MEG) would not be an option due to capsule ferromagnetic components, though as previously mentioned, one MEG study detailed evidence of midline parieto-occipital activity linked to stomach input under resting conditions (30). Several unanswered questions remain. For example, what are the molecular and cellular entities transducing the delivered stimulation? And what are the associated peripheral (presumably neural) pathways conveying the mechanosensory vibration signals to the brain? Answering these questions would likely require the application of invasive (i.e., nonhuman) studies capable of evaluating mechanotransducers at molecular levels. One candidate would be PIEZO channels (13, 14), which are heavily expressed in the stomach (51, 52) – though many other mechanical (e.g., transient receptor potential (TRP) channels) and voltage-gated ion channels are possibilities (either individually or in combination). At the cellular level, several types of mechanosensitive neurons within the enteric nervous system are known to play a role in transmitting tensile and mechanical forces, including extrinsic vagal afferent nerve endings in the upper gut and Dogiel type II neurons (spinal afferent nerve endings, which predominantly innervate the lower gut might be considered less likely)(53). Stomach stretching also elicits mechanosensory signaling that is relayed via GLP1R neurons (a vagal afferent subtype) to autonomic brainstem nuclei (nucleus tractus solitarius and area postrema)(54), providing a plausible pathway by which vibratory stimulation might reach the brain. Speculation as to the subsequent trafficking of signals in the brain is beyond the scope of the current study, though others have pointed to the role of hierarchical homeostatic reflexes in the transmission (17) and regulation (55) of information across the brain-gut loop. These feedback loops are certainly amenable to perturbation in a variety of contexts using the gut-brain probe demonstrated here.

In conclusion, a minimally invasive mechanosensory probe of stomach sensation elicits intensity dependent increases in perceptual accuracy and event related potentials in parieto-occipital EEG leads. These changes are significantly associated with one another and are unrelated to myoelectric indices of gastric rhythm. This finding of a gastric evoked potential provides an opportunity to better understand the role of gastrointestinal symptoms in a variety of human pathological conditions and may ultimately provide insights into how gut feelings are processed by the human brain.

## Materials and Methods

### Participants

Healthy adult male and female volunteers between the ages of 18 to 40 years were recruited from the general Tulsa community through electronic and print advertisements. Eligibility was verified via completion of structured medical and psychiatric screening evaluations. Exclusion criteria included current pregnancy or positive for drugs of abuse as defined by a urine screen during screening and during the stimulation visit, current diagnosis of psychiatric disorders, history or current diagnosis of a significant gastrointestinal disorder, gastrointestinal surgery, or other medical disorder involving respiratory, cardiovascular, renal, hepatic, biliary or endocrine disease, as well as chronic use of psychotropic medications or non-steroidal anti-inflammatory drugs. The study was conducted at the Laureate Institute for Brain Research and the research protocol was approved by the Western Institutional Review Board (IRB). All participants provided written informed consent and received financial compensation for participation.

### Vibrating Capsule

The vibrating capsule was developed by Vibrant Ltd and is under investigation as a non-pharmacologic therapeutic option for chronic constipation via delivery of stimulation in the colon. It consists of an orally administered non-biodegradable capsule that is wirelessly activated using an activation base unit (Figure 1A). The Vibrant capsule is a non-significant risk device (NSR). The safety of this approach has been established in healthy human volunteers (56) and in patients with chronic constipation (57, 58).

### Masking Procedure

To constrain expectancies participants were told that two different modes of the Vibrant capsule were being evaluated and that they would be randomly assigned to one of three arms of the study: capsule mode A or B (during which the capsule would vibrate), or a placebo capsule that did not vibrate. Participants were further informed that neither they nor the experimenter would know whether any stimulations will occur. However, in actuality, every participant received a capsule that delivered vibratory stimulations, making this a single-blinded protocol. To ensure that the contents of the stomach were empty at the time of capsule ingestion participants were instructed to begin fasting (defined as no food or drink for 3 hours prior to the study visit).

### Mechanosensory Stimulation

Delivery of mechanosensory stimulations to the stomach started shortly following ingestion of the Vibrant capsule. Capsule activation occurred by placing the capsule in the base unit. Shortly after activation participants ingested the capsule with approximately 240 milliliters of water while seated in a chair. They were subsequently asked to attend to their stomach sensations while resting their eyes on a fixation cross displayed on a monitor approximately 60 centimeters away. They were instructed to use their dominant hand to press and hold a button each time they felt a sensation that they ascribed to the capsule and to release the button as soon as this sensation had ended (Figure 1B). Stimulations began approximately 3 minutes after capsule activation in the base unit. Participants remained seated throughout the experiment in order to reduce movement artifact in the EEG and EGG signals. They were asked to rest their non-dominant hand in their lap as well as to avoid palpating their abdomen. They were visually observed throughout the experiment by a research assistant seated behind them to verify alertness and compliance with these instructions.

In the experiment, each participant received two blocks of vibratory stimulation (normal and enhanced) in a counterbalanced order. The normal condition entailed the delivery of a standard level of mechanosensory stimulation (as developed by Vibrant) matching the level of stimulation delivered during chronic constipation trials targeting the colon. The enhanced condition entailed the delivery of an increased level of mechanosensory stimulation, which was expected to facilitate gastrointestinal perception. Each block included a total of 60 stimulations (each 3 seconds in duration), which were delivered in a pseudorandom order across a 13-minute period. After a 4-minute pause, a second round of 60 stimulations were delivered in pseudorandom order during a 13-minute period. Thus, participants rated the presence of gastrointestinal sensations throughout a 33-minute period following capsule ingestion. This timing ensured the capsule remained in the stomach during stimulations since the normal gastric emptying time is estimated to be approximately 30 minutes (59). Due to a technical, error 3 vibrations were missing from the enhanced vibration block in the normal/enhanced sequence.

### Vibration Detection

To precisely verify the vibration timing a digital stethoscope (Thinklabs Inc.) was gently secured against the anterior surface of the lower right quadrant of the abdomen using a Tegaderm patch (15 × 20 centimeters). The associated signal was continuously recorded during the entire experiment and fed into the physiological recording software at a sampling rate of 1000 Hz. To identify vibrations, we developed custom analysis scripts in Matlab (Mathworks, Inc.) and used a two-step procedure to detect the onset and offset of each delivered vibration. In the first step, the script detected the vibration timings automatically using the “*findchangepts*” function in Matlab. In the second step, the timing graph for each vibration was visually inspected and adjusted if needed. Participants for whom the amplitude of vibrations could not be confidently identified using this procedure were excluded from analysis (2 individuals; 1 had a faulty capsule that did not vibrate and was also excluded). An example of the stethoscope-derived vibration waveforms for the enhanced and normal vibration conditions are shown in supplementary figure S5.

### Electrophysiological Recordings

EEG signals were recorded continuously using a 32-channel EEG system from Brain Products GmbH, Munich, Germany. The EEG cap consisted of 32 channels, including references, arranged according to the international 10-20 system (Figure 1C). One of these channels recorded the electrocardiogram (ECG) signal via electrode placement on the back, leaving 31 EEG signals available for analysis. The EEG signal was acquired at a sampling rate of 5000 Hz and a measurement resolution of 0.1 microvolts (μV).

Besides EEG, three additional physiological measures were used in this study, including electrogastrogram (EGG), skin conductance response (SCR), and electrocardiogram (ECG). All three physiological measurements were acquired using a Biopac MP150 Acquisition Unit (Goleta, California) with a sampling frequency of 1000 Hz. Acquisition of EEG and psychological measures was synchronized via a TTL pulse signal continuously fed from the EEG/button response computer to the peripheral physiological recording computer. Cutaneous EGG signals were captured via two abdominal electrodes positioned below the left costal margin and between the xiphoid process and umbilicus. The reference electrode was positioned in the right upper quadrant in line with a previous study (60) (Figure 1C). SCRs were recorded using gel-filled electrodes attached to the thenar and hypothenar eminences of the nondominant palm. The ECG was recorded using two electrodes positioned in a lead-2 placement. All recordings were screened for physiological artifacts (e.g., motion) and analyzed offline using AcqKnowledge 3.5.

### EEG Data Processing

The pre/post-processing of EEG data was completed using BrainVision Analyzer 2 software (Brain Products GmbH, Munich, Germany). EEG data was downsampled to 250 Hz. Next, a fourth-order Butterworth (i.e., 24 dB/octave roll-off) band-rejection filter (1 Hz bandwidth) was used to remove alternating current (AC) power line noise (60 Hz). Then, a bandpass filter between 0.1 and 80 Hz (eighth order Butterworth Filter, 48 dB/octave roll-off) was utilized to remove signals unrelated to brain activity. Afterward, the infomax independent component analysis (ICA) was applied for independent component decomposition (61) over the entire data length after excluding intervals with excessive motion-artifact. ICA was run on the data from 31 EEG channels yielding 31 independent components (ICs). The time course signal, topographic map, power spectrum density, and energy of these ICs were utilized to detect and remove artifactual ICs (i.e., ocular, muscle and single channel artifacts) (62). In addition to these preprocessing steps, the data was segmented from the 200 milliseconds (ms) prior to the 3000 ms post onset of each vibration. Then the data were baseline corrected to the average of the 200 ms interval preceding the vibration onset. EEG data was referenced to the average of the mastoid channels (TP9 and TP10). Finally, automated procedures were used to detect bad intervals and flatlining in the data. Bad intervals were defined as any change in amplitude between data points that exceeded 50μv or absolute fluctuations exceeding 200μV in any 200 ms interval of the segments (i.e., −200 to 3000 ms); flat lining was defined as any change of less than 0.5μV in a 200 ms period. Trials were excluded if they included any of these artifacts.

### Peripheral Physiological Data Processing

#### Electrogastrogram (EGG)

The single-channel EGG recording from each subject was divided into three blocks: baseline, normal stimulation, and enhanced stimulation based on the counterbalanced protocol. For each block, the spectral power was computed to identify the location with the largest activity in the normogastria range (2.5-3.5 cycle per minute (cpm)). The spectral power analysis retained peaks of frequency in each condition for each subject. Fast Fourier Transform (FFT) from the FieldTrip toolbox (63) was used to estimate the spectral power with a Hanning taper to reduce spectral leakage and control frequency smoothing. To further characterize the gastric rhythm, we adopted a finite impulse response (FIR) filter to filter the EGG signal into low-frequency ranges. FIR copes very well with very low-frequency filtering (as shown in (64)). Then, we applied a Hilbert transform to compute the instantaneous phase and amplitude envelope of the gastric rhythm. To further account for bad segments in the data, we adopted the artifact detection method described in (64). This method relies on the regularity of the computed cycle durations (the standard deviation (STD) of cycle duration from the condition). More specifically, a segment was considered as an artifact if the cycle length was greater than the mean ± STD of the cycle length distribution or if the cycle showed a nonmonotonic change in phase. Following the decision tree approach, any cycle with either of these conditions was considered as bad interval and excluded from the signal. The power spectral analysis was calculated again after excluding bad segments from the EGG signal, including subsequent filtering. We report the absolute power for four gastric ranges: normogastria [2.5-3.5] cpm, tachygastria [3.75 –9.75] cpm, bradygastria [0.5 –2.5] cpm, and total power [0.5 –11] cpm (in line with (65)).

#### Heart Rate (HR)

To estimate the HR changes for phasic stimulatioon (during vibration period; 3 secs) and pre-stimulation segments (no vibration period; preceding 3 secs), the peak of the R-waves was detected using a custom peak detection algorithm in MATLAB. For each block, the difference in HR between the duration of the stimulus (3 sec) and pre-stimulus (3 sec) period was reported. In order to have a baseline comparison, we added 60 pseudo-events with 30-sec intervals into the 30 minute baseline period (after ignoring the first 2 minutes to allow for reaching a physiological steady state). Additionally, we estimated the overall HR for each block as the average HR across 60-sec windows.

#### Heart Rate Variability (HRV)

R-wave peaks were used to calculate the heart rate variability for each block. We extracted the Interbeat intervals (IBIs) for each block independently. Then a custom IBI outlier detection implemented in the HRVAS Toolbox in MATLAB (66) was used to remove abnormal IBIs. After that, the standard deviation of the normal (NN) sinus-initiated IBI (SDNN) was calculated for each block.

#### Skin Conductance Response (SCR)

We used Continuous Deconvolution Analysis (CDA) implemented in the Ledalab Toolbox (67) to characterize phasic changes in SCRs between stimulation and pre-stimulation segments. Specifically, we used the maximum value of phasic activity with a threshold of 0.01 muS to quantify SCRs in each segment. Prior to phasic changes extraction, we downsampled the signal to 20 Hz and applied a smoothing window of 200. Similar to the HR analysis, we used the same pseudo-events from the baseline period to characterize changes in SCR relative to physiological resting conditions.

#### Breathing Rate (BR)

We did not use a breathing belt to minimize the delivery of skin-mediated feedback during capsule vibrations, so we estimated BR from the ECG signal. To do so, we adapted a custom MATLAB script to estimate the BR using the R-R interval peaks via an autocorrelation approach to estimate the periodic maxima corresponding to integer multiples of the signal’s fundamental frequency (68). To improve algorithmic estimation of BR we divided each block into 60-second windows while applying detrending of the ECG signal resulting in 27, 13, and 13 windows for the baseline, normal, and enhanced blocks, respectively. Then, we estimated BR for each block as the average across all windows.

### Subjective Rating Measures

Participants completed several self-report surveys and visual analog scales indexing subjective and metacognitive experiences related to capsule stimulation including the perceived intensity of stomach/digestive, breath, heartbeat, and muscle sensations. The scale for the intensity ratings ranged from 0 (“Not at all/None”) to 100 (“Extremely/The most I have ever felt”). Trait scales were completed during the screening visit, with the remainder of the state scales completed before and after the capsule stimulation.

### Perceptual Discrimination Measures

We calculated measures of interoceptive accuracy for each block and participant, adopting a non-parametric signal detection analog of d′ used for conditions with low trial numbers (69, 70) as follows:

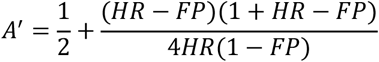

where HR represents the Hit Rate and FP is the False positive rate. We further normalized the A′ (A prime) scores using the following equation: 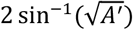, resulting in values between 0 and π. In addition, we calculated the average and the STD of the response latency for each block and each participant. Response latency was defined as the difference between the vibration onset and the participant’s button press response indicating a perceived sensation. Each participant’s performance was further evaluated using a binomial test to check if they performed above chance during the normal and enhanced blocks, defined as ≥ 70 correct trials (out of a possible total of 120, counting the 60 normal vibration and 60 nonvibration intervals as trials) and ≥ 67 correct trials (out of a possible total of 114, counting the 57 enhanced vibration and 57 nonvibration intervals as trials) corresponding to an individual performance threshold statistically above chance (p < 0.05) according to the binomial distribution (as in (70)). *Note:* One participant did not generate any button press responses during the normal block; therefore, there was no average and STD response latency available for this individual, and no line connecting their performance from the normal to enhanced blocks for those measures (e.g., in Figure 2B).

### Statistical Analysis

Two-way analyses of variance (ANOVA) with the block (normal, enhanced) and sex (male, female) as independent variables and separated subject-level performance measures (normalized A’, of sex and block in each of those perception measures. 2-way repeated measure ANOVAs were also utilized to examine the differences between EEG channels (i.e., CP1, CP2, P3, Pz, P4, POz, O1, Oz, and O2) and blocks (normal, enhanced), EGG power in different frequency bands (normogastria, tachygastria, bradygastria) and different blocks (baseline, normal, and enhanced), as well as comparing the subjective measure of 4 different intensities (i.e., stomach/digestive, breath, heartbeat, and muscle) and time (pre-, post-stimulus). For the repeated measures ANOVA, if the assumption of sphericity was violated according to Mauchly’s test for any variables, degrees of freedom were adjusted using Greenhouse-Geisser estimates of sphericity. We applied 1-way repeated measure of ANOVA for comparing the total EGG power, SCR, tonic HR, phasic HR, HRV and phasic BR in 3 different blocks (baseline, normal, enhanced), with post-hoc testing using two-tailed t tests following the observation of significant main effects. Several paired t tests (two-tailed) were conducted for comparing the behavioral measurements between then normal and enhanced blocks. The nonparametric Spearman’s rank correlation was used to evaluate the association between neurophysiological measurements and accuracy measurements due to the non-normal distributions of such measurements. The statistical analyses were performed using SPSS version 26 (IBM SPSS Inc., Chicago, Ill).

## Data and code availability

Access to the data collected using the Vibrant capsule is covered by nondisclosure and material transfer agreements between LIBR and Vibrant Ltd. The raw and/or processed data may be made available upon request to researchers, pending the establishment of similar agreements between all parties.

## Acknowledgments

We would like to thank Dhvanit Raval, Chloe Sigman, Katie Baker, Ann Marie Flusche, and Maria Puhl for assistance with data pre-processing, Valerie Upshaw for assistance with data collection, and Gurpreet Singh and Jennifer Stewart for helpful discussions. A portion of Figure 1 was created with BioRender. Funding for the study was provided by The William K. Warren Foundation.

## Abbreviations

BR: Breathing Rate
CPM: Cycles per minute
EEG: Electroencephalogram
EGG: Electrogastrogram
ERP: Evoked response potential
FP: False positive
HR: Hit rate
HRV: Heart Rate Variability
LPP: Late positive potential
M: Mean
μV: Microvolts
SCR: Skin Conductance Response
STAI: Spielberger State Trait Anxiety Inventory
STD: Standard deviation.

## Notes

**Competing Interest Statement:** The authors declare that they have no competing interests.

### Competing Interest Statement

The authors have declared no competing interest.

