## Supplement for "Neural indicators of human gut feelings" (from another version of this preprint: version 1, 2021-02-13)

***Supplemental information***

Ahmad Mayeli[1], Obada Al Zoubi[1], Evan J. White[1], Sheridan Chappelle[1], Rayus Kuplicki[1], Ryan Smith[1],

Justin S. Feinstein[1], Jerzy Bodurka[1,2], Martin P. Paulus[1,3], Sahib S. Khalsa[1,3]*

[1]Laureate Institute for Brain Research, Tulsa, OK, United States
[2]Stephenson School of Biomedical Engineering, University of Oklahoma, Tulsa, OK, United States
[3]Oxley College of Health Sciences, University of Tulsa, Tulsa, OK, United States

*** Correspondence:**
Sahib S. Khalsa
6655 S Yale Ave
Tulsa, OK 74136-3326


**This PDF file includes:**

Figures S1 to S2
Tables S1 to S6

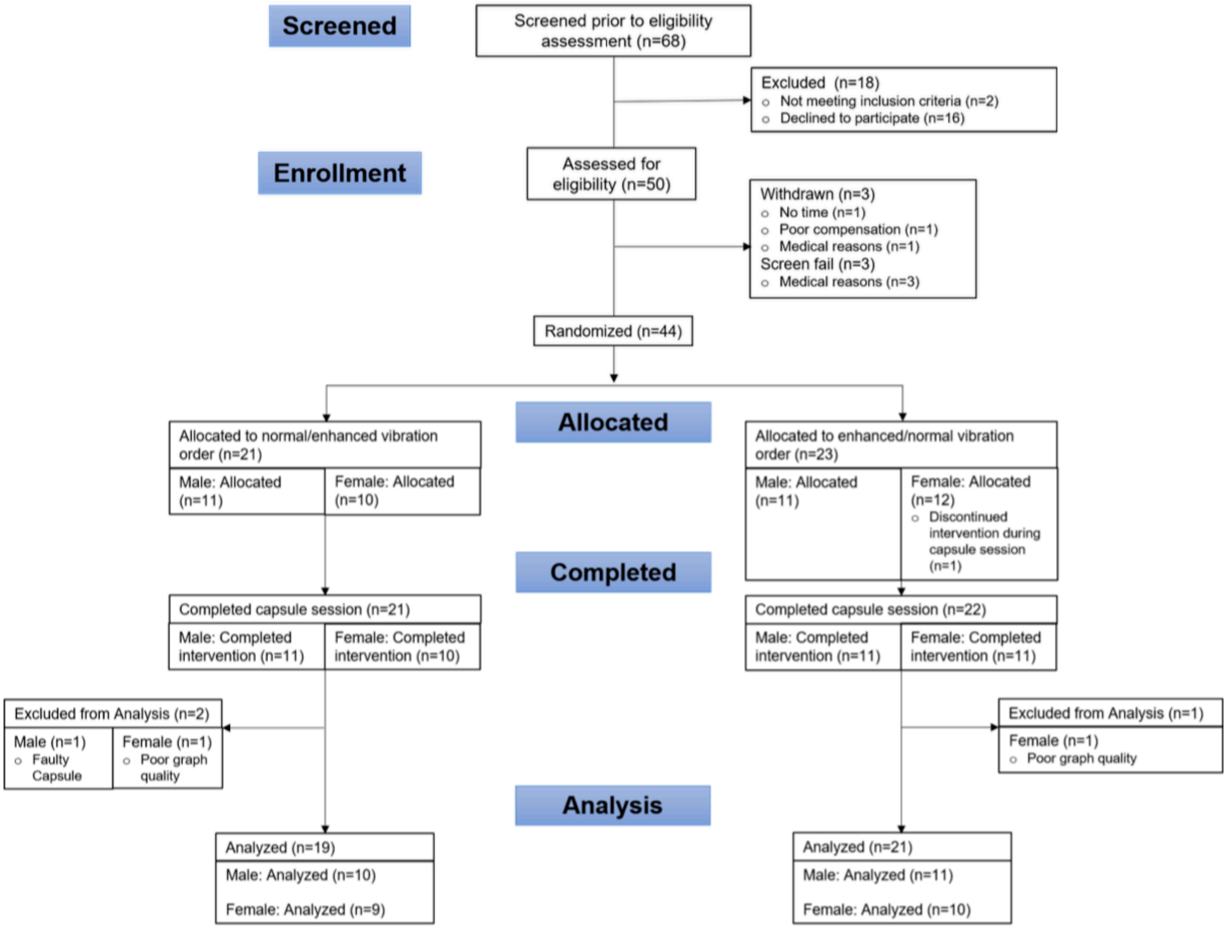


**Fig. S1**: Consort diagram showing the flow of study participants.

**Table S1:** Separate general linear model results comparing the Normalized A Prime, Average Response Latency, and the Standard Deviation (STD) of Response Latency, with Sex and Block (i.e., normal and enhanced) as fixed factors.

| | *Normalized A Prime* | | *Average Response Latency* | | *STD Response Latency* | |
|---|---|---|---|---|---|---|
| | F-Value | P-Value | F-Value | P-Value | F-Value | P-Value |
| *Sex* | 0.000334 | 0.985 | 0.081 | 0.777 | 0.055 | 0.816 |
| *Block* | 22.008 | 0.000012 | 25.032 | 0.000004 | 26.111 | 0.000002 |
| *Sex*Block* | 0.141 | 0.708 | 0.008 | 0.93 | 0.193 | 0.662 |

**Table S2:** Estimated marginal ERP means for the parieto-occipital electrode sites between 300 ms and 600 ms.

| Channel | Mean | Std. Error | 95% Confidence Interval | |
|---|---|---|---|---|
| | | | **Lower Bound** | **Upper Bound** |
| **P3** | 2.591 | 0.403 | 1.762 | 3.420 |
| **P4** | 2.957 | 0.454 | 2.024 | 3.891 |
| **Cz** | 1.406 | 0.422 | 0.537 | 2.274 |
| **Oz** | 2.536 | 0.387 | 1.741 | 3.330 |
| **POz** | 3.746 | 0.482 | 2.755 | 4.737 |
| **Pz** | 3.849 | 0.505 | 2.812 | 4.886 |

**Table S3:** Pairwise comparison of ERP means for the parieto-occipital electrode sites between 300 ms and 600 ms.

| **(I) Channel** | **(J) Channel** | **Mean Difference (I-J)** | **Std. Error** | **Sig.[b]** | **95% Confidence Interval for Difference[b]** | |
|---|---|---|---|---|---|---|
| | | | | | Lower Bound | Upper Bound |
| **P3** | P4 | -0.366 | 0.194 | 1.000 | -0.993 | 0.261 |
| | Cz | 1.186 | 0.442 | 0.189 | -0.244 | 2.615 |
| | Oz | 0.055 | 0.305 | 1.000 | -0.931 | 1.042 |
| | POz | -1.155 | 0.182 | 0.000* | -1.743 | -0.567 |
| | Pz | -1.258 | 0.184 | 0.000* | -1.852 | -0.663 |
| **P4** | P3 | 0.366 | 0.194 | 1.000 | -0.261 | 0.993 |
| | Cz | 1.552 | 0.445 | 0.026* | 0.115 | 2.988 |
| | Oz | 0.421 | 0.359 | 1.000 | -0.739 | 1.581 |
| | POz | -.789 | 0.169 | 0.001 | -1.335 | -0.243 |
| | Pz | -.892 | 0.155 | 0.000* | -1.393 | -0.390 |
| **Cz** | P3 | -1.186 | 0.442 | 0.189 | -2.615 | 0.244 |
| | P4 | -1.552 | 0.445 | 0.026* | -2.988 | -0.115 |
| | Oz | -1.130 | 0.600 | 1.000 | -3.070 | 0.809 |
| | POz | -2.340 | 0.524 | 0.002* | -4.032 | -0.649 |
| | Pz | -2.443 | 0.412 | 0.000* | -3.774 | -1.112 |
| **Oz** | P3 | -0.055 | 0.305 | 1.000 | -1.042 | 0.931 |
| | P4 | -0.421 | 0.359 | 1.000 | -1.581 | 0.739 |
| | Cz | 1.130 | 0.600 | 1.000 | -0.809 | 3.070 |
| | POz | -1.210 | 0.291 | 0.005* | -2.151 | -0.269 |
| | Pz | -1.313 | 0.426 | 0.072 | -2.689 | 0.063 |
| **POz** | P3 | 1.155 | 0.182 | 0.000* | 0.567 | 1.743 |
| | P4 | .789 | 0.169 | 0.001* | 0.243 | 1.335 |
| | Cz | 2.340 | 0.524 | 0.002* | 0.649 | 4.032 |
| | Oz | 1.210 | 0.291 | 0.005* | 0.269 | 2.151 |
| | Pz | -0.103 | 0.207 | 1.000 | -0.773 | 0.567 |
| **Pz** | P3 | 1.258 | 0.184 | 0.000* | 0.663 | 1.852 |
| | P4 | .892 | 0.155 | 0.000* | 0.390 | 1.393 |
| | Cz | 2.443 | 0.412 | 0.000* | 1.112 | 3.774 |
| | Oz | 1.313 | 0.426 | 0.072 | -0.063 | 2.689 |
| | POz | 0.103 | 0.207 | 1.000 | -0.567 | 0.773 |

[b] Adjustment for multiple comparisons: Bonferroni. *indicates significant electrode site contrast.

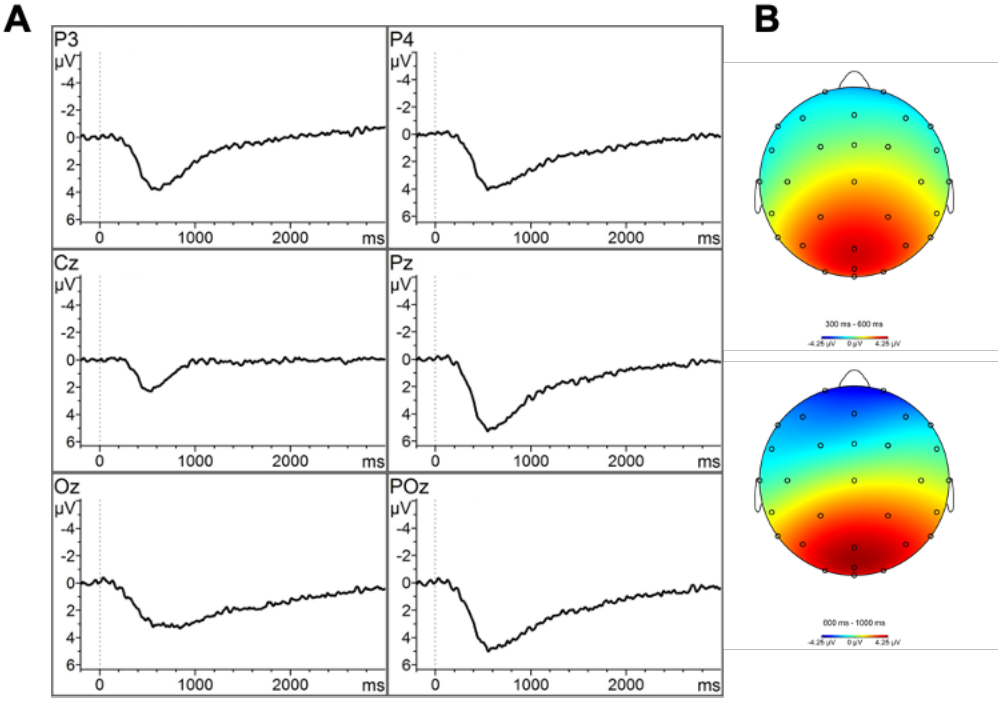


**Fig. S2: A)** ERP waveforms for channels P3, P4, Cz, Pz, Oz, and POz across the whole experiment, obtained by combining the normal and enhanced blocks. Time-zero represents the onset of the capsule vibration. Waveform averages were calculated from the mastoid reference electrode. **B)** Late positive potential (LPP) topography across the scalp at two windows: between 300 and 600 ms after vibration onset (relative to the 200 ms pre-stimulation baseline) and between 600 and 1000 ms relative to baseline. Note: These results only include subjects with >50% correct responses during vibrations (n=38) in order to maximize power for the ERP analysis.

**Table S4:** Estimated marginal means for the interoceptive intensity ratings. The rating scale ranged from 0 (“Not at all/None”) to 100 (“Extremely/The most I have ever felt”).

| **Measurement** | **Time** | **Mean** | **Std. Error** | **95% Confidence Interval** | |
|---|---|---|---|---|---|
| | | | | Lower Bound | Upper Bound |
| **Stomach/Digestive** | Pre- stimulation | 23.55 | 2.149 | 7.729 | 16.421 |
| | Post- stimulation | 54.22 | 3.346 | 16.183 | 29.717 |
| **Breath** | Pre- stimulation | 12.07 | 2.079 | 4.669 | 13.081 |
| | Post- stimulation | 22.95 | 2.472 | 10.451 | 20.449 |
| **Heartbeat** | Pre- stimulation | 8.875 | 2.954 | 17.574 | 29.526 |
| | Post- stimulation | 15.45 | 3.572 | 47 | 61.45 |
| **Muscle** | Pre- stimulation | 17.25 | 2.329 | 12.54 | 21.96 |
| | Post- stimulation | 14.35 | 2.039 | 10.226 | 18.474 |

**Table S5:** Separate general linear model results comparing the stomach/digestive, breath, heartbeat, and muscle with sex and time (i.e., pre- and post- stimulation) as fixed factors.

| | *Stomach/Digestive* | | *Breath* | | *Heartbeat* | | *Muscle* | |
|---|---|---|---|---|---|---|---|---|
| | F-Value | P-Value | F-Value | P-Value | F-Value | P-Value | F-Value | P-Value |
| *Sex* | 0.046 | 0.830 | 0.184 | 0.669 | 0.308 | 0.581 | 2.365 | 0.128 |
| *Block* | 43.705 | 4.69E-9 | 8.627 | 0.004 | 4.422 | 0.039 | 0.751 | 0.389 |
| *Sex*Block* | 0.665 | 0.417 | 3.489 | 0.066 | 0.795 | 0.375 | 0.497 | 0.483 |

**Table S6:** Time and events checklist for the screening and stimulation visits.

| **Procedures** | Visit 1 Screening Enrollment | Visit 2 Before Simulation | Visit 2 After Stimulation |
|---|---|---|---|
| Informed consent | X | | |
| Demographics | X | | |
| Medical history | X | | |
| Current medications and interim medical procedures | X | X | |
| Vital Signs | X | X | X |
| Height (Meters) | X | | |
| Weight (Kilograms) | X | | |
| Urine drug and pregnancy screen[a] | X | X | |
| Randomization | X | | |
| State Trait Anxiety Inventory | X | X | X |
| Interoceptive VAS ratings | | X | X |

[a]Women of childbearing potential
